# Helitrons are enriched in lichenized fungi with long generation lengths and small distribution sizes

**DOI:** 10.1101/2025.07.11.664412

**Authors:** Julianna Paulsen, Stephen T. Sharrett, Devin Mumey, Elaine M. Larsen, Nguyen Khoi Nguyen, James Lendemer, Lalita M. Calabria, Jordan R. Hoffman, Krisztian Magori, Jessica L. Allen

**Affiliations:** Department of Biology, Eastern Washington University, Cheney, WA 99004, U.S.A.; Department of Botany, Research & Collections, The New York State Museum, Albany, NY 12230, U.S.A.; The Evergreen State College, 2700 Evergreen Parkway NW, Olympia, WA 98505, U.S.A.; Division of Science and Mathematics, Delta College, University Center, MI 48710, U.S.A.; NatureServe, Arlington, VA 22202, U.S.A

## Abstract

Mobile genetic elements (MGEs) have the potential to drive genome evolution by introducing mutations and causing structural instability and chromosomal rearrangements, particularly under conditions like environmental or genetic stress. In this study, we generated 18 new long-read based reference genomes for lichenized fungi, which form obligate mutualistic symbioses with algae or cyanobacteria. We used the new genomes and 10 publically available genomes to investigate the relationships between species traits (i.e., dominant reproductive mode, distribution size, and generation length) and the abundance and spatial distribution of MGEs using a phylogenetic comparative framework. We found that species with smaller distribution sizes and longer generation lengths had a higher genomic DNA transposon load. Specifically, their genomes were enriched with Rolling Circle transposons, which contradicts previous research that has identified high proportions of retrotransposons in rare species. Disproportionate distributions of MGEs in rare and range-restricted species may disrupt genomic stability, decrease fitness, and be reflective of species experiencing a greater degree of stress. Conversely, greater MGE activity may be an important source of novel genetic diversity in isolated populations with limited gene flow. Further research is needed to understand the potential mechanisms driving MGE proliferation in rare species’ genomes, and if MGE content is predictive of increased extinction risk.

## INTRODUCTION

Mobile genetic elements (MGEs) are genes that encode their own machinery to move within a genome (Wells and Feschotte 2020). In Eukaryotes, MGEs are divided into two classes. Class I MGEs, called retrotransposons or retroelements (RTs), are distinguished by the use of an mRNA intermediate to complete transposition, and are often described as ‘copy and paste’ elements. Class II elements, called DNA transposons or transposable elements (TEs), do not use an mRNA intermediate, and instead transpose directly through a mechanism described as ‘cut-and-paste.’ TEs are generally not self-replicative, but are capable of utilizing host machinery to increase their copy number in a genome. One mechanism by which TE proliferation occurs is during the S phase of mitosis, where TEs can move from replicated to unreplicated chromosomes, duplicating themselves in the process (Zaratiegui 2017). MGEs are subclassified by the specifics of their genetic structure, such as variations in the genes and proteins involved in these processes, insertion site preferences, and the presence of accessory genes (Wicker et al. 2007). Helitrons, for example, are a type of TE that transpose via a rolling-circle mechanism using the RepHel protein, enabling them to self-replicate during each transposition event (Barro-Trastoy and Köhler 2024). Subclass characteristics affect the transposition efficiency, replication, and selective pressures related to host fitness (Duhamel et al. 2023; Bourgeois et al. 2020).

MGEs have the potential to facilitate genomic restructuring and evolution by reorganizing genes and chromosomes and causing nucleotide mutations (Almojil et al. 2021; Rahnama et al. 2020; Wong et al. 2019; Kent et al. 2017; Aksenova et al. 2013; de Jonge et al. 2013). The activity level of MGEs is an important determinant of their proliferation efficiency and, subsequently, their interactions with the genome. Interactions between host defenses and MGE offenses are often described as an evolutionary arms race, where both parties engage in a rapid diversification of strategies to outpace the other (Koonin et al. 2020). One such strategy employed by eukaryotic hosts to curb the potentially deleterious effects of MGE transposition includes silencing of MGEs by specialized RNA molecules, which recruit DNA methylation machinery or bind to transcripts directly and trigger cleavage (Lax et al. 2020). That being said, MGE success in eukaryotes is intrinsically tied to host fitness, as MGEs must proliferate in the germline to persist within a population. Thus, MGEs self-regulate their expression using encoded transcriptional regulators and modifying their insertion preferences (Bonnet and Lesage 2021; Hermant and Torres-Padilla 2021). MGE activity is known to be positively correlated with stress, and it is believed that cellular stress responses divert resources away from quelling MGEs, allowing their proliferation in the genome (Fouché et al. 2019; Mustafin and Khusnutdinova 2019).

Previous studies have reported high genomic MGE loads in rare and endemic species across kingdoms (Wang et al. 2023; Linscott et al. 2022; Ruiz-Ruano et al. 2021), but what drives this pattern is not known. Environmental conditions and population structure are traits correlated with rarity that may be connected to MGE abundance in the genome (Ruiz-Ruano et al. 2021; Abascal et al. 2016). Additionally, certain classes of MGEs, such as RTs, may be correlated with adverse environmental conditions to which rare species are often subjected (Milyeava et al. 2023; Cavrak et al. 2014). A better understanding of the relationship between rarity and MGEs is pertinent for conservation efforts, as large scale alterations of the genome facilitated by MGEs can decrease fitness and increase the rate of speciation (Ricci et al. 2018; Serrato-Capuchina and Matute 2018). In the case of endangered organisms, either outcome translates to increased extinction risk.

Lichens are obligate mutualistic symbioses composed of dominant fungal, algal, and/or cyanobacterial symbionts, and a variety of associated microbes and macrobes. They exhibit diverse life history traits that underlie their interactions with the environment, such as different reproductive strategies, substrate preferences, growth forms, and symbiotic associations (Manzitto-Tripp et al. 2022). Due to their relevance as quintessential symbioses and their ability to produce diverse and biotechnologically relevant secondary metabolites, lichen genomics has become a popular avenue of research (Singh et al. 2025; Allen and Lendemer 2022). However, their MGE landscapes remain less well characterized than other taxonomic groups. In the small number of published, long-read genomes of lichenized fungi where MGE content has been reported, RTs are identified as the most abundant MGE, and total MGE content ranges from 8-33% (Heuberger et al. 2024; Allen et al. 2021; Mckenzie et al. 2020). This is in line with fungal genomes more broadly, that are reported to range from 0-45%, the majority of which are also identified as RTs (Castanera et al. 2017).

In this study, we investigate the relationship between MGEs and rarity, reproductive mode, generation length, and distribution size in lichenized fungi using a multi-species, comparative genomics framework. Using genomes generated via long read sequencing for 26 different species of North American Lecanoromycete lichenized fungi, 18 of which represent previously unpublished sequences, we identified MGEs, calculated multiple metrics for rarity, and divided them into groups based on rarity-associated life history traits. We hypothesized that MGEs, specifically retrotransposons, would be present in greater abundance in the genomes of rare species. We tested this hypothesis by comparing the number of MGEs and the percent of the genome occupied by MGEs for each species in a phylogenetically informed statistical framework. Additionally, we compared the spatial distribution of MGEs, hypothesizing that rare species would experience higher instances of colocalization of genes and MGEs. To better understand intraspecific variation in the species with the highest MGE content, we compared the MGE abundance and genomic structural variation between two *Pseudocyphellaria rainierensis* individuals from different populations. We found that DNA transposon abundance was significantly associated with distribution size and generation length. We also found that Helitrons were enriched in the genomes of species with small distribution sizes and long generation lengths.

## METHODS

### Genome Generation and Annotation

We generated 18 new genomes in this study (Table 1). Genomes were sequenced and analyzed using standard lab procedures published in Pfeffer et al. (2023). After collection, specimens were air dried then either frozen until extraction or extractions were initiated within 24 hours of collection. Specimens were divided into 16 tissue disruption tubes, and DNA extraction was performed using the DNeasy Plant Pro Kit (Qiagen, Germany) according to the manufacturer’s protocol with minor modifications. TissueLyser II (Qiagen, Germany) was used to disrupt dry material, and this step was repeated until the specimen was thoroughly homogenized. Resultant DNA was eluted using 50 microliters of elution buffer, and the final extracts were pooled into a single tube per species. Any remaining tissue after DNA extraction was deposited in the Eastern Washington University herbarium. DNA was quantified with the Qubit Fluorometer 4.0 using the 2x HSdsDNA reagents following the manufacturer’s protocol (Thermo Fisher Scientific, United States). To select for high-molecular weight DNA fragments a 0.4:1 ratio of Mag-Bind TotalPure NGS magnetic beads (Omega BioTek, United States) to extract was used for size selection in all specimens except *Sticta beauvoisii*, where a 0.8:1 ratio was used due to initially low DNA yield. Sequencing was completed using the Nanopore MinION Mk1B (Nanopore, United Kingdom). Oxford Nanopore Technologies Ligation sequencing kit (LSK-114) library preparations were completed according to the manufacturer’s protocol for the majority of genomes (Nanopore, United Kingdom). Genome sequencing for *Lepraria finkii*, *L. lanata*, *L. normandinoides*, *L. oxybapha*, *Punctelia appalachensis*, *P. rudecta*, *Usnea strigosa*, and *Usnea subfusca* were completed by Cold Spring Harbor Laboratories on the Oxford Nanopore Technologies PromethION platform. Basecalling was completed with Dorado v0.9.6 (Nanopore, United Kingdom), and assembly of metagenomic data was completed using Flye v2.9.5 in meta mode (Kolmogorov et al. 2020). Metagenomes were filtered based on the workflow described in McKenzie et al. (2020), which relies on identifying the dominant fungal symbiont through sorting contigs based on depth, GC content, and homology searches. Genomes were annotated for repetitive regions using RepeatModeler2 v2.0.6 (Flynn et al. 2020) and Repeatmasker v4.1.8 (Smit et al. 2015). All other annotations were conducted using AntiSMASH v6.1.1 (Blin et al. 2025) and Funannotate v1.8.7 for cleaning, masking, and predicting, and Funannotate v1.8.17 for final annotations (Palmer and Stajich 2020).

**Table 1.** Genome Metrics for newly generated genomes, previously published internal genomes, and external genomes acquired from GenBank.

| Species | Total<br>Genome<br>Size | Number of<br>Contigs | N50 | GC<br>Content | Number of<br>Genes | Number of<br>tRNAs | Complete<br>BUSCOs | Single<br>BUSCOs | Duplicate<br>BUSCOs |
| --- | --- | --- | --- | --- | --- | --- | --- | --- | --- |
| *Cladonia chlorophaea* | 42625121 | 523 | 148102 | 45.29 | 12952 | 99 | 71.5 | 71.5 | 0 |
| *Cladonia imbricarica* | 36666740 | 37 | 1603632 | 42.92 | 8954 | 60 | 92.8 | 92.7 | 0.1 |
| *Cladonia rangiferina* | 38484624 | 36 | 1569438 | 42.98 | 9923 | 57 | 93.5 | 93.4 | 0.1 |
| *Diploschistes muscorum* | 51546914 | 111 | 1537671 | 41.23 | 8401 | 108 | 91.7 | 91.4 | 0.3 |
| *Lepraria finkii* | 40852740 | 32 | 4154807 | 46.58 | 9409 | 84 | 94.1 | 94 | 0.1 |
| *Lepraria lanata* | 47102205 | 42 | 2355529 | 45.6 | 9775 | 62 | 94.8 | 94.6 | 0.2 |
| *Lepraria normandinoides* | 29300421 | 162 | 403430 | 48.41 | 7927 | 137 | 91.8 | 91.8 | 0 |
| *Lepraria oxybapha* | 4.34E+07 | 109 | 1210317 | 46.18 | 9144 | 82 | 91.8 | 91.8 | 0 |
| *Pseudocyphellaria citrina* | 51635776 | 44 | 2349121 | 47.17 | 13698 | 84 | 94.4 | 94.4 | 0 |
| *Pseudocyphellaria*<br>*rainierensis (Surp. Creek)* | 125582678 | 26 | 7148722 | 39.02 | 13824 | 64 | 93.9 | 93.9 | 0 |
| *Pseudocyphellaria*<br>*rainierensis (Col. Bob)* | 131031989 | 19 | 7147418 | 39.13 | 14480 | 46 | 90.9 | 90.9 | 0 |
| *Punctelia appalachensis* | 45689027 | 25 | 2208748 | 40.97 | 9991 | 65 | 94.8 | 94.6 | 0.2 |
| *Punctelia rudecta* | 42749237 | 31 | 1874871 | 43.77 | 11857 | 58 | 94.9 | 94.7 | 0.2 |
| *Sticta beauvoisii* | 46674901 | 184 | 677788 | 47.71 | 10458 | 57 | 94.1 | 94 | 0.1 |
| *Sticta deyana* | 43509696 | 127 | 891764 | 48.13 | 10601 | 60 | 93.9 | 93.9 | 0 |
| *Sulcaria isidiifera* | 71390556 | 23 | 3455340 | 32.4 | 10429 | 91 | 95.2 | 95 | 0.2 |
| *Usnea strigosa* | 43172064 | 32 | 2168579 | 42.49 | 10511 | 89 | 90.1 | 89.9 | 0.2 |
| *Usnea subfusca* | 49089431 | 30 | 2173746 | 41.37 | 10855 | 64 | 94.8 | 94.7 | 0.1 |
| Previously Published Genomes (internal) |  |  |  |  |  |  |  |  |  |
| *Bacidia gigantensis* | 33117897 | 24 | 1807239 | 44.67 | 8942 | 51 | 90.7 | 90.5 | 0.2 |
| *Lepraria neglecta* | 42023604 | 11 | 5201525 | 47.39 | 11669 | 43 | 95 | 95 | 0 |
| *Letharia columbiana* | 5.23E+07 | 161 | 666803 | 39.57 | 11564 | 76 | 91.9 | 91.8 | 0.1 |
| *Letharia lupina* | 49242005 | 31 | 2098233 | 38.73 | 10214 | 60 | 95.5 | 95.4 | 0.1 |
| Externally Published Genomes |  |  |  |  |  |  |  |  |  |
| *Acarospora socialis* | 31172923 | 38 | 2402664 | 47.3 | 8587 | 109 | 97.4 | 97.4 | 0 |
| *Bacidia rubella* | 33524925 | 51 | 1771855 | 45.28 | 9589 | 45 | 93.8 | 93.6 | 0.2 |
| *Cladonia perforata* | 33606327 | 72 | 1102137 | 47.06 | 8624 | 46 | 95.5 | 95.4 | 0.1 |
| *Niebla homalea* | 50566743 | 52 | 1266640 | 37.96 | 9409 | 67 | 93.6 | 93.4 | 0.2 |
| *Ramalina farinacea* | 32742913 | 44 | 1546935 | 46.69 | 9087 | 64 | 93.9 | 93.6 | 0.3 |
| *Umbilicaria muhlenbergii* | 34812353 | 7 | 7009248 | 47.12 | 8148 | 35 | 96.5 | 96.5 | 0 |

### Comparative Genomics Analysis

Comparative genomics analyses were performed using 18 newly generated genomes, four genomes previously published from our research group (Pfeffer et al. 2023; Allen et al. 2021; McKenzie et al. 2020) and six publically available genomes acquired from GenBank (Table 2).

**Table 2.** Species Metadata including collection info for new genomes, source publication for previously published genomes, trait data used in this study, and sources for distribution sizes and trait data for each species.

| Species Name | Collection Metadata | Distribution Size (km2) | Generation Length | Reproductive Mode | Rarity | Trait Data Sources |
| --- | --- | --- | --- | --- | --- | --- |
| *Cladonia chlorophaea* | *Pinus ponderosa* savanna, Palisades park, Rimrock Trail, Washington, Spokane Co. (47.656 N, -117.484 W; Devin Mumey 420) | 13725112.52 | short | A | C | Tripp and Lendemer (2020) |
| *Cladonia imbricarica* | *Pinus ponderosa* savanna, Dishman Hills Conservation Area, just west of Iller Creek Trailhead, Washington, Spokane Co. (47.602 N, -117.283 W; Devin Mumey 407-A) | 1035578.519 | short | A | R | Brodo et al. (2001) |
| *Cladonia rangiferina* | Exposed soil outcrop embankment along infrequently traveled roadside. Washington, Chelan Co., Leavenworth, Wenatchee Nat Forest, Roadside of NF-6701 past intersection with NF-6700, between the bridge over Rainy Creek and other river further along the road. (47.8595 N, -120.9835 W; Jordan Hoffman s.n.) | 12778158.25 | short | A | C | Tripp and Lendemer (2020) |
| *Diploschistes muscorum* | Turnbull Lab for Ecological Studies, 0.12 mi NE of turnoff of S. Badger Lake Road, 1.5 mi S of intersection between Lt. Col. Michael P. Anderson Memorial Highway and Cheney Plaza Road. Washington, Spokane Co., Cheney. (47.45481 N, -117.5744 W; Jessica Allen 5013) | 14407856.16 | long | S | C | Tripp and Lendemer (2020) |
| *Lepraria finkii* | Great Smoky Mountains National Park, Lumber Ridge Trail ~2.3 mi NE of Tremont Institute, S slopes of Lumber Ridge above Bloody Branch of Meigs Creek. Mixed hardwood (*Acer saccharum*, *Carya*, *Oxydendrum*, *Quercus alba*, *Q. rubra*) - conifer (*Pinus*, *Tsuga*) forest with *Kalmia-Rhododendron* understory. On *Quercus rubra* base. (35.6531 N, - 83.6711 W; James Lendemer 71953) | 11214498.23 | short | A | C | Tripp and Lendemer (2020) |
| *Lepraria lanata* | NW-facing overhang on Cattail Peak, North Carolina (35.7964 N, - 82.2569 W; Allen s.n.) | 28112.47664 | medium | A | R | Tripp and Lendemer (2020) |
| *Lepraria normandinoides* | vertical shale outcrop at the pigeon River Whitewater Boat Launch, Tennessee (35.7756N, - 83.101 W; Hollinger 27,155) | 1361504.869 | short | A | C | Tripp and Lendemer (2020) |
| *Lepraria oxybapha* | quartzite outcrop in Great Smoky National Park, on a roadbank off Old Cataloochee Turnpike, North Carolina (35.6465 N, -83.0633 W; Hollinger 27,129) | 1369424.654 | short | A | C | Tripp and Lendemer (2020) |
| *Pseudocyphellaria citrina* | United States of America, Washington State, Alpine Lakes Wilderness, Taylor River Trail, FS trail 1002. (47.57408 N, -121.42597 W; Gifford (Marco) Pinochet IV s.n.) | 787956.7691 | long | A | C | Lücking et al. (2017) |
| *Pseudocyphellaria rainierensis* (Col. Bob) | *Tsuga heterophylla* - *Abies amabilis* / *Polystichum*<br>*munitum-Vaccinium ovalifolium* stand with varying levels of riparian influence. Moderate to dense coarse woody debris. Understory sparse/patchy with lots of *Rubus spectabilis* along riparian areas. Colonel Bob Trail #851, near intersection with Ziegler Creek, 3 mi from TH, Colonel Bob Wilderness, Olympic National Forest, Washington (47.47786 N, - 123.763650 W; Stephen Sharrett 105) | 259693.6853 | long | A | R | Lücking et al. (2017) |
| *Pseudocyphellaria rainierensis* (Surp. Creek) | *Tsuga-Thuja plicata* climax stand with scattered *Abies amabilis* and riparian influence. Moderate to dense coarse woody debris. Lots of recent windfall. Open understory - dominated by *Vaccinium ovalifolium*/*Cornus canadensis*/*Clintonia uniflora*. Lots of small seeps/stream running south lined with *Rubus*<br>*spectabilis* / *Oplopanax horridus*. Surprise Creek. NE of the Surprise Creek Trail #1060. Approx. 0.42 mi S of the Surprise Creek Trail TH, 4.51 mi SSW of Steven's Pass. Alpine Lakes Wilderness, Washington. (47.7018 N, -121.15705 W; Stephen Sharrett 903) | 259693.6853 | long | A | R | Lücking et al. (2017) |
| *Punctelia appalachensis* | Cataloochee divide, Great Smoky Mountain National Park, North Carolina (35.585 N, -83.074 W; Allen 6097) | 187104.6771 | medium | A | R | Tripp and Lendemer (2020) |
| *Punctelia rudecta* | Cataloochee divide, Great Smoky Mountain National Park, North Carolina (35.585 N, -83.074 W; Allen 6098) | 3587472.473 | medium | A | C | Tripp and Lendemer (2020) |
| *Sticta beauvoisii* | Metasandstone outcrop, Locust Run Creek, Virginia (38.7240 N, -77.9273 W; Harris s.n.) | 544060.7335 | medium | A | C | Brodo et al. (2001) |
| *Sticta deyana* | Sipsey wilderness area, Alabama (34.3241 N, -87.4513 W; Tripp 6599) | 1552.481037 | long | A | R | Tripp and Lendemer (2020) |
| *Sulcaria isidiifera* | Elfin Forest, California (35.3325 N, 120.8238 W; | 36.19660427 | long | A | R | McMullin et al. (2019) |

|  | Balderas 70) |  |  |  |  |  |
| --- | --- | --- | --- | --- | --- | --- |
| *Usnea strigosa* | Cataloochee divide, Great Smoky Mountain National Park, North Carolina ( 35.582012, N, 14 83.07037; Allen 6099) | 2003165.168 | medium | S | C | Tripp and Lendemer (2020) |
| *Usnea subfusca* | West slopes of Jack Mountain, Highlands Wildlife Management Area, Virginia (38.3746, - 79.5034; Allen 6100) | 48785.13062 | medium | S | R | Tripp and Lendemer (2020) |
| Previously Published Genomes |  |  |  |  |  |  |
| Species Name | Genome Source Citation |  |  |  |  | Sources |
| *Bacidia gigantensis* | Allen et al. (2021) | 209.1420958 | medium | S | R | Allen et al. (2021) |
| *Lepraria neglecta* | Pfeffer et al. (2023) | 10426807.23 | short | A | C | Tripp and Lendemer (2020) |
| *Letharia columbiana* | McKenzie et al. (2020) | 1202898.439 | medium | S | C | Altermann et al. (2016) |
| *Letharia lupina* | McKenzie et al. (2020) | 2395797.465 | medium | A | C | Altermann et al. (2016) |
| *Acarospora socialis* | Adams et al. (2023) | 6249824.495 | long | S | C | Nash et al. (2002) |
| *Bacidia rubella* | Gerasimova et al. (2022) | 1152705.446 | medium | S | C | Tripp and Lendemer (2020) |
| *Cladonia perforata* | Leavitt et al. (2023) | 12123.31399 | short | A | R | Tripp and Lendemer (2020) |
| *Niebla homalea* | Duong et al. (2021) | 43307.77313 | long | S | R | Brodo et al. (2001) |
| *Ramalina farinacea* | Llewellyn et al. (2023) | 1414099.471 | medium | A | C | Brodo et al. (2001) |
| *Umbilicaria muhlenbergii* | Park et al. (2014) | 4686307.09 | long | S | C | Tripp and<br>Lendemer (2020) |

The dataset contains a total of 28 genomes representing 27 different species. Two genomes of the species *Pseudocyphellaria rainierensis* were generated for individuals from different populations (Colonel Bob Wilderness, Washington, USA (Col. Bob); Surprise Creek, Washington, USA (Surp. Creek)) to investigate intraspecific variation in genomic MGE content. One genome, *Cladonia chlorophaea*, was not sufficiently complete and was thus not included in any statistical analyses. All previously published genomes were reannotated according to methods above to ensure standardized comparisons across genomes. We first sought to build a robust phylogeny to place all of our analyses in a comparative phylogenetic framework. To build the phylogeny, we used a total 50 genomes of Lecanoromycete lichenized fungi and closely related ascomycetes, utilizing *Saccharomyces cerevisieae* and *Schizosaccharomyces pombe* as outgroups. This included the 28 genomes in our dataset, twenty-one additional genomes generated with short-read data originating from Gerasimova et al. (2022), and the long-read reference genome for *Pseudevernia furfuracea* (GCA_022315425.1) acquired from GenBank. Orthofinder v2.5.5 was then used to infer phylogenetic relationships using orthologous genes (Emms and Kelly 2019).

Phytools v2.4.4 was used to load and configure the species tree in R for subsequent analyses (Revell 2011). Because contiguous repetitive sequences cannot be as accurately assembled with short-read data or be spatially localized in reference to genes across the genome, we trimmed the species tree to only include long-read reference genomes for subsequent analyses.

Trait data, including rarity (rare or common), reproductive mode (sexual or asexual), generation length (short, medium, or long), and distribution size (meters squared) were compiled from expert knowledge, publications, and federal data (Table 2). Rarity was assigned by holistically assessing various life history traits, including those analyzed in this study, as well as additional factors like patchiness, abundance, and changes in population sizes that can be evaluated by field observations but are not easily collapsible into a discrete categorical or numerical trait. Therefore, rarity as it is used here captures more ecological nuance than any one single trait alone, allowing us to evaluate whether specific traits or combinations thereof are more influential of MGE constituency. Generation length was defined based on methods from Yahr et al. (2024), where species are grouped into categories of generation length based on their life history strategies. Distribution sizes were calculated in ArcGIS v3.30 (ESRI 2024) by georeferencing expert drawn distribution maps or referencing published species descriptions (Table 2), transforming them into shapefiles, clipping the polygons to the North American continental landmass, and erasing all terrestrial water features using shapefiles from the Commission for Environmental Cooperation (Montreal, CA; 2023 and 2022). To improve the accuracy of area calculation, geospatial data was projected onto an Albers equal area conic basemap. The final area of each polygon was reported in square meters using the calculate geometry tool, and this data was exported using a custom Python script.

Because related species tend to be more similar in terms of genetic traits than unrelated ones, we tested for phylogenetic signal in our statistical models using the R package phyr v1.1.0 (Li et al. 2020). The presence of statistically significant phylogenetic signal in the models justified the use of phylogenetically informed statistical tests for downstream analyses (Supplementary Table 1). We tested for correlations among trait data using non-parametric statistical methods: Chi-square tests for associations between two-level categorical traits, Wilcoxon rank-sum tests for comparisons between continuous variables and two-level categorical traits, and Kruskal-Wallis tests for comparisons involving continuous variables and three-level categorical traits. Our analyses revealed a significant relationship between only rarity and log-transformed distribution size (Supplementary Table 2). Univariate phylogenetic generalized linear mixed models (PGLMM) from the package phyr v1.1.0 (Li et al. 2020) were used to test the relationships between both continuous and discrete trait data and both metrics of MGE constituency, number of elements and percent of genome occupied. To assess if RTs are enriched in the genomes of rare species in relation to TEs, we calculated the percentage of total MGEs that were RTs for each species and conducted PGLMM analyses on the resulting proportions. Distribution size was log transformed for all analyses due to the presence of extreme low and high values. We corrected for increased family-wise type I error using a Bonferroni correction within each set of analyses.

To take an in-depth look at the role of MGEs in intraspecific variation, we aligned the genomes of two individuals of *Pseudocyphellaria rainierensis* collected from different populations and assessed the resulting alignment for the presence of structural variants (SVs) caused by MGEs. To serve as a comparison, the same analysis was conducted on one closely related species pair in the dataset, *Sticta beauvoisii* and *Sticta deyana* (Supplementary Figure 1). Minimap2 v2.28 (Li 2018) was used to create the alignment, Sniffles2 v2.6.0 (Smolka et al. 2024) was used to identify and visualize SVs, and D-Genies v1.5.0 (Cabanettes and Klopp 2018) was used to visualize the alignments.

### MGE Spatial Distribution Analysis

To determine if genes and MGEs tend to colocalize more often in rare species than common ones, we analyzed the spatial distribution of MGEs relative to genes. General Feature Format (GFF3) files containing the annotations generated by Funannotate (Palmer and Stajich 2020) and RepeatModeler2 (Flynn et al. 2020) were loaded into R as GRanges objects using the R package GenomicRanges v1.54.1 (Lawrence et al. 2013) and ape v5.8 (Paradis and Schliep 2018). We calculated the distance between each MGE and the nearest gene on the same contig, and summarized the mean of this value for each species. This analysis was then repeated after subsetting the data by strand, such that the distances were calculated between MGEs and the nearest gene on the same strand and in the direction of transcription to capture MGE association with both primary and quaternary genomic structures, where transcription levels can vary (Figure 1a). Mean distances were corrected for differences in genome size and gene density by multiplying the values by the total number of genes by total base pair per genome. To assess the relationship between average MGE to gene distances and trait data we used a univariate Gaussian PGLMM.

**Fig 1.**
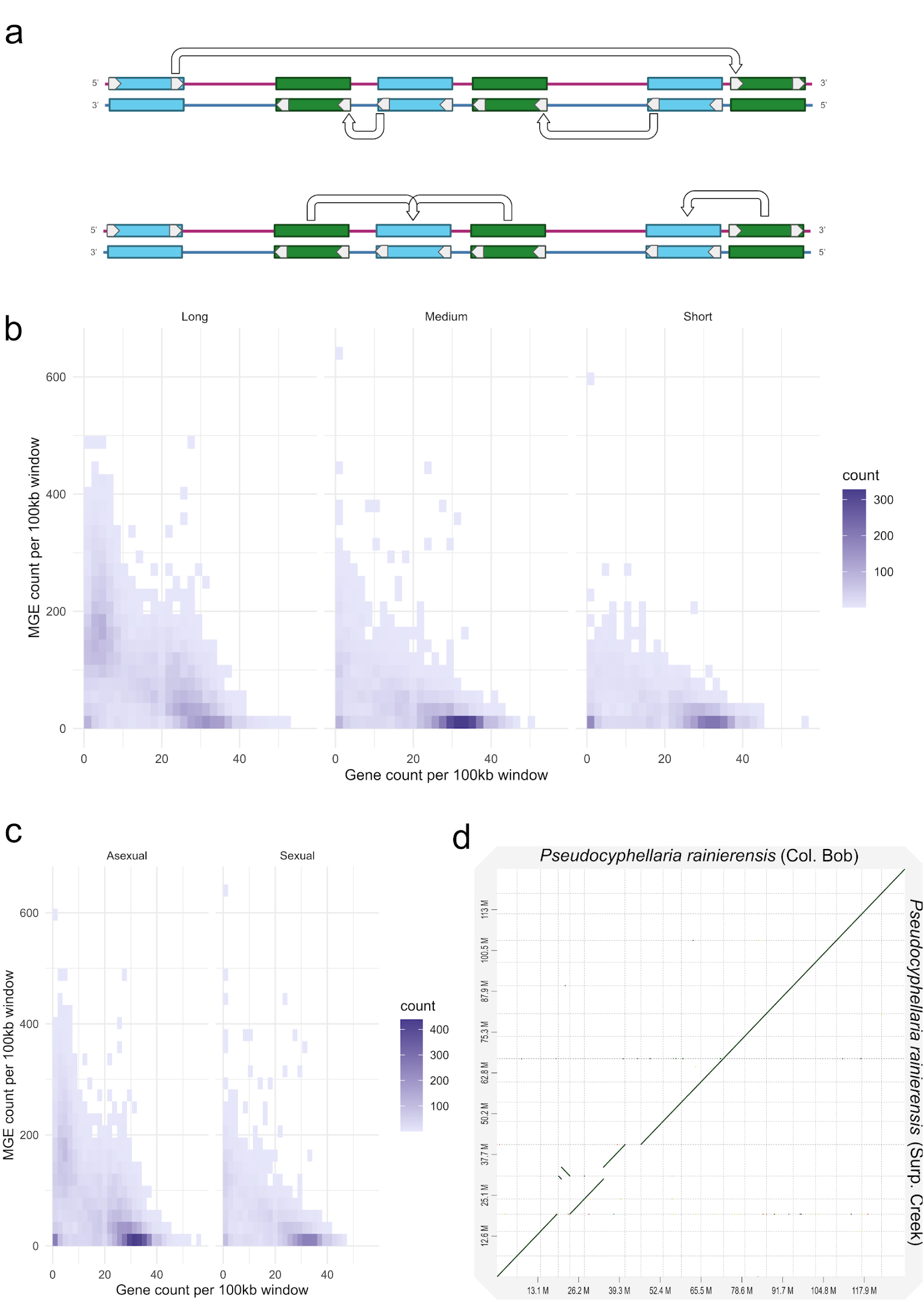
MGE impacts on genome structure. **a)** Schematic representation of the method to calculate mobile genetic element (MGE) gene colocalization. The first analysis (top) was conducted by calculating the distance between the end of each gene (blue) and the start of the nearest MGE (green) on the same strand, in the direction of transcription, and within the same contig. The second analysis was conducted by calculating the distance between the midpoint of each MGE and the nearest gene, regardless of strand and direction. **b)** Heatmap of the number of 100kb windows in each genome (represented by the gradient of low values (light purple) to high values (dark purple)) with a specific number of genes (x-axis) and MGEs (y-axis) by generation length. If differences in MGE gene colocalization were observed, it would be expected that high values of the color gradient would be present in different quadrants of the graphs for each group. **c)** The same gene-MGE distance heatmap described in (b) faceted by reproductive mode. **d)** Dotplot of the alignment of two *Pseudocyphellaria rainierensis* genomes collected and sequenced from two different populations. Gridlines delineate contigs. Total genome length is represented on the bottom and left axes.

## RESULTS

### Genome Metrics

Eighteen genomes were newly generated as part of this study resulting in a total of 1.2 Gb of data after assembly and filtering (Table 1). The average N50 value was 2.4 Mb. The highest N50 was 7.15 Mb for *Pseudocyphellaria rainierensis* (Surp. Creek), and the lowest value was 148.1 Kb belonging to *Cladonia chlorophaea*. Genome assembly completeness assessments using BUSCO v2.0 (Manni et al. 2021) recovered a high number of conserved genes in all genomes (>90%) except *Cladonia chlorophaea,* which was dropped from downstream analyses because the low BUSCO score indicated that the assembly was incomplete and may therefore underrepresent MGE constituency. We added 10 published genomes generated with long-read sequence data to our dataset. Thus, our final dataset for the MGE analysis contained 27 genomes from 26 different species of lichenized fungi. The complete set of 27 genomes was annotated with the same exact pipeline to ensure comparable results. A total of 289,022 genes were annotated across all 28 genomes. *Pseudocyphellaria rainierensis* (Col. Bob) had the highest number of genes annotated at 14,480, and *Lepraria normandinoides* had the lowest number of genes annotated at 7,927.

RepeatModeler2 recovered MGEs in all genomes in the analysis (Table 3). The mean percent of the genome occupied by MGEs was generally higher in rare species than in common ones for TEs, RTs, and total MGEs which remained consistent for the number of MGEs identified and the total length of MGEs identified in base pairs (Figure 3a, b; Table 3). The percent of the genome occupied by MGEs ranged from 4.6% in *Bacidia gigantensis* to 69.55% in *Pseudocyphellaria rainierensis*, averaging 23.48% for all species. For TEs, the percent of genome occupied ranged from 0% in *Bacidia gigantensis* and *Cladonia rangiferina* to 19.88% in *Sulcaria isidiifera,* averaging 4.38% among all species. Retrotransposons ranged from 3.9% in *Sticta deyana* to 65.49% in *Pseudocyphellaria rainierensis*, averaging 18.86% across all species.

**Table 3.**
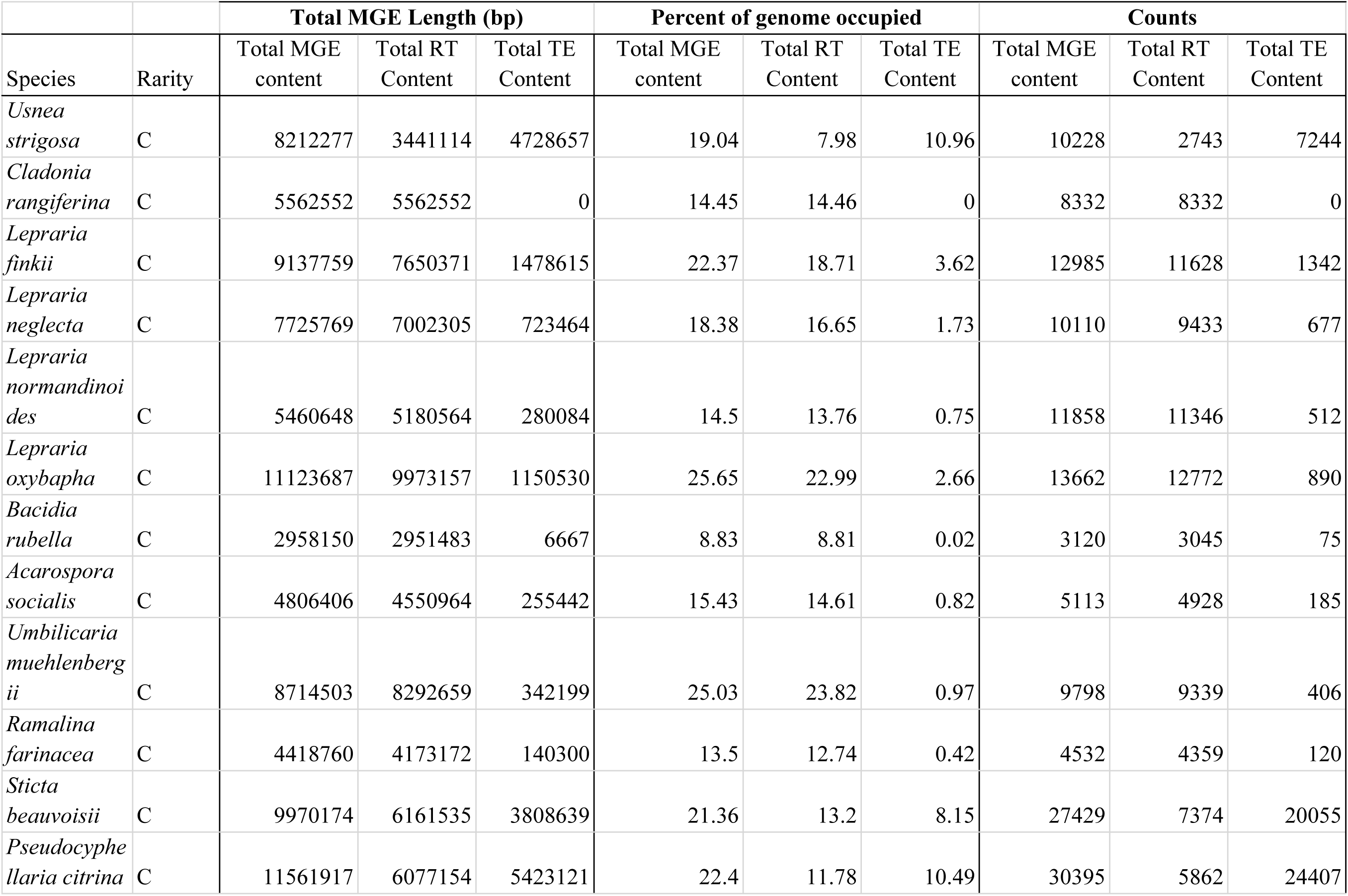
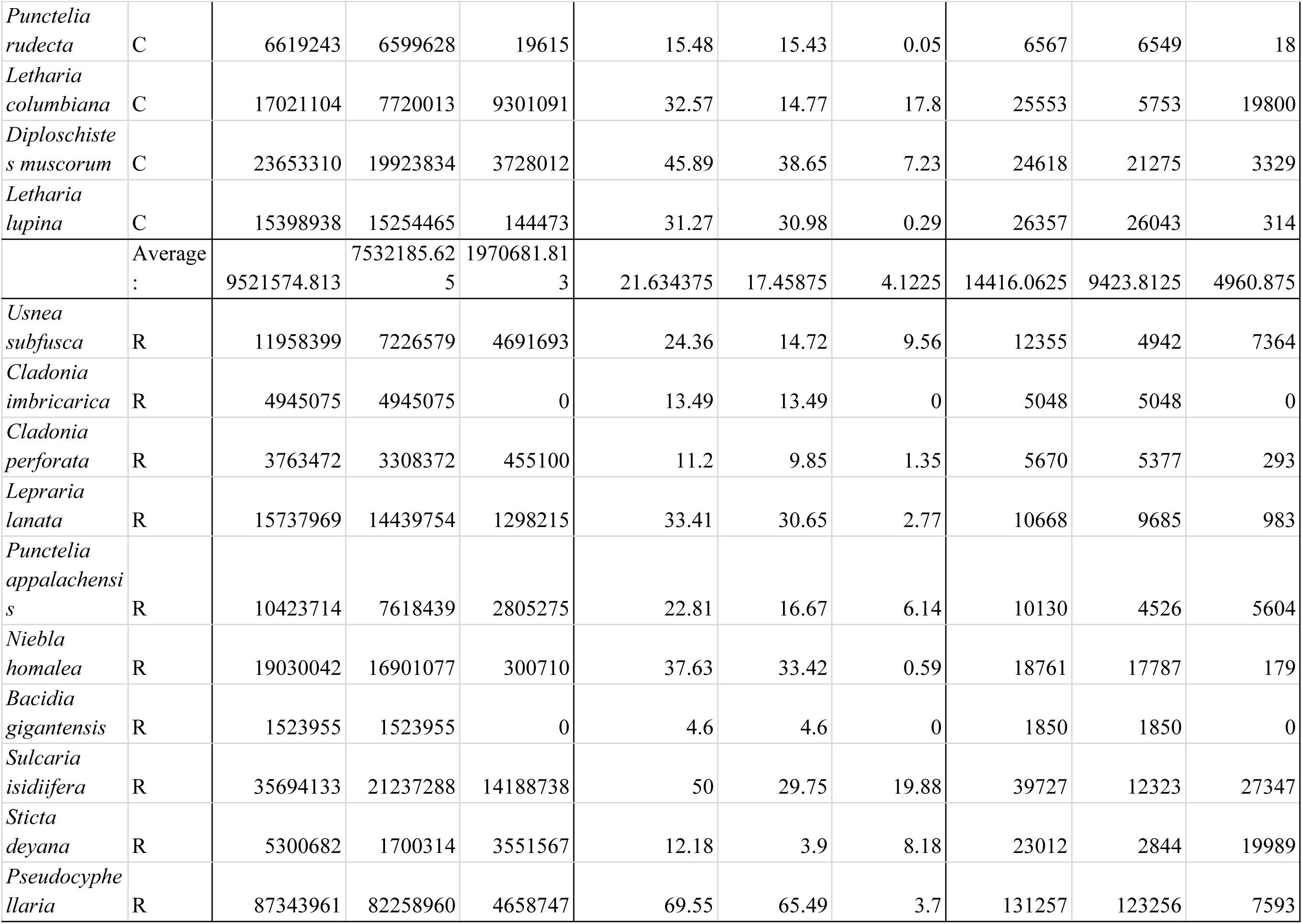
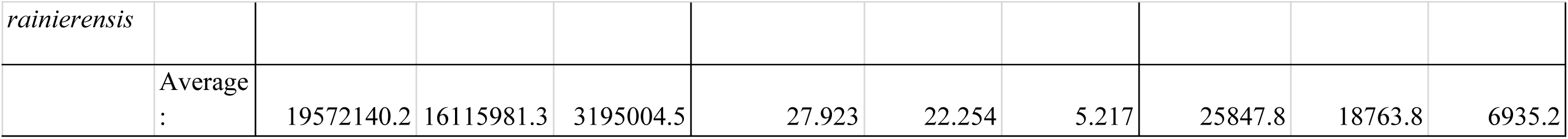
Summary of Mobile Genetic Element (MGE) Results for percent of genome occupied, MGE counts, and bp length by rarity.

| Species | Rarity | Total MGE Length (bp) |  |  | Percent of genome occupied |  |  | Counts |  |  |
| --- | --- | --- | --- | --- | --- | --- | --- | --- | --- | --- |
|  |  | Total MGE content | Total RT Content | Total TE Content | Total MGE content | Total RT Content | Total TE Content | Total MGE content | Total RT Content | Total TE Content |
| *Usnea strigosa* | C | 8212277 | 3441114 | 4728657 | 19.04 | 7.98 | 10.96 | 10228 | 2743 | 7244 |
| *Cladonia rangiferina* | C | 5562552 | 5562552 | 0 | 14.45 | 14.46 | 0 | 8332 | 8332 | 0 |
| *Lepraria finkii* | C | 9137759 | 7650371 | 1478615 | 22.37 | 18.71 | 3.62 | 12985 | 11628 | 1342 |
| *Lepraria neglecta* | C | 7725769 | 7002305 | 723464 | 18.38 | 16.65 | 1.73 | 10110 | 9433 | 677 |
| *Lepraria normandinoi des* | C | 5460648 | 5180564 | 280084 | 14.5 | 13.76 | 0.75 | 11858 | 11346 | 512 |
| *Lepraria oxybapha* | C | 11123687 | 9973157 | 1150530 | 25.65 | 22.99 | 2.66 | 13662 | 12772 | 890 |
| *Bacidia rubella* | C | 2958150 | 2951483 | 6667 | 8.83 | 8.81 | 0.02 | 3120 | 3045 | 75 |
| *Acarospora socialis* | C | 4806406 | 4550964 | 255442 | 15.43 | 14.61 | 0.82 | 5113 | 4928 | 185 |
| *Umbilicaria muehlenbergii* | C | 8714503 | 8292659 | 342199 | 25.03 | 23.82 | 0.97 | 9798 | 9339 | 406 |
| *Ramalina farinacea* | C | 4418760 | 4173172 | 140300 | 13.5 | 12.74 | 0.42 | 4532 | 4359 | 120 |
| *Sticta beauvoisii* | C | 9970174 | 6161535 | 3808639 | 21.36 | 13.2 | 8.15 | 27429 | 7374 | 20055 |
| *Pseudocyphe llaria citrina* | C | 11561917 | 6077154 | 5423121 | 22.4 | 11.78 | 10.49 | 30395 | 5862 | 24407 |
| *Punctelia rudecta* | C | 6619243 | 6599628 | 19615 | 15.48 | 15.43 | 0.05 | 6567 | 6549 | 18 |
| *Letharia columbiana* | C | 17021104 | 7720013 | 9301091 | 32.57 | 14.77 | 17.8 | 25553 | 5753 | 19800 |
| *Diploschistes muscorum* | C | 23653310 | 19923834 | 3728012 | 45.89 | 38.65 | 7.23 | 24618 | 21275 | 3329 |
| *Letharia lupina* | C | 15398938 | 15254465 | 144473 | 31.27 | 30.98 | 0.29 | 26357 | 26043 | 314 |
|  | Average : | 9521574.813 | 7532185.625 | 1970681.813 | 21.634375 | 17.45875 | 4.1225 | 14416.0625 | 9423.8125 | 4960.875 |
| *Usnea subfusca* | R | 11958399 | 7226579 | 4691693 | 24.36 | 14.72 | 9.56 | 12355 | 4942 | 7364 |
| *Cladonia imbricaria* | R | 4945075 | 4945075 | 0 | 13.49 | 13.49 | 0 | 5048 | 5048 | 0 |
| *Cladonia perforata* | R | 3763472 | 3308372 | 455100 | 11.2 | 9.85 | 1.35 | 5670 | 5377 | 293 |
| *Lepraria lanata* | R | 15737969 | 14439754 | 1298215 | 33.41 | 30.65 | 2.77 | 10668 | 9685 | 983 |
| *Punctelia appalachensis* | R | 10423714 | 7618439 | 2805275 | 22.81 | 16.67 | 6.14 | 10130 | 4526 | 5604 |
| *Niebla homalea* | R | 19030042 | 16901077 | 300710 | 37.63 | 33.42 | 0.59 | 18761 | 17787 | 179 |
| *Bacidia gigantensis* | R | 1523955 | 1523955 | 0 | 4.6 | 4.6 | 0 | 1850 | 1850 | 0 |
| *Sulcaria isidiifera* | R | 35694133 | 21237288 | 14188738 | 50 | 29.75 | 19.88 | 39727 | 12323 | 27347 |
| *Sticta deyana* | R | 5300682 | 1700314 | 3551567 | 12.18 | 3.9 | 8.18 | 23012 | 2844 | 19989 |
| *Pseudocyphellaria* | R | 87343961 | 82258960 | 4658747 | 69.55 | 65.49 | 3.7 | 131257 | 123256 | 7593 |
| *rainierensis* |  |  |  |  |  |  |  |  |  |  |
|  | Average<br>: | 19572140.2 | 16115981.3 | 3195004.5 | 27.923 | 22.254 | 5.217 | 25847.8 | 18763.8 | 6935.2 |

### Phylogenetic analyses and trait data

To generate a robust phylogeny for downstream phylogenetically-informed statistical analyses, we used whole genome protein sequence annotations from our focal 26 species along with a suite of publicly available fungal genomes generated with short-read data (Supplementary Figure 2). Orthofinder uncovered a total of 593,001 genes among the 58 genomes used to generate the species tree, of which 570,340 (96%) were within the 25,506 orthogroups recovered. Three-hundred ninety-six of the orthologues were present in all species in the analysis. All branch support values on the resulting species tree were 100%, except the placement of *Cyanodermella asteris*. All nodes in the final pruned tree had support values of 100% (Figure 2a).

**Fig 2.**
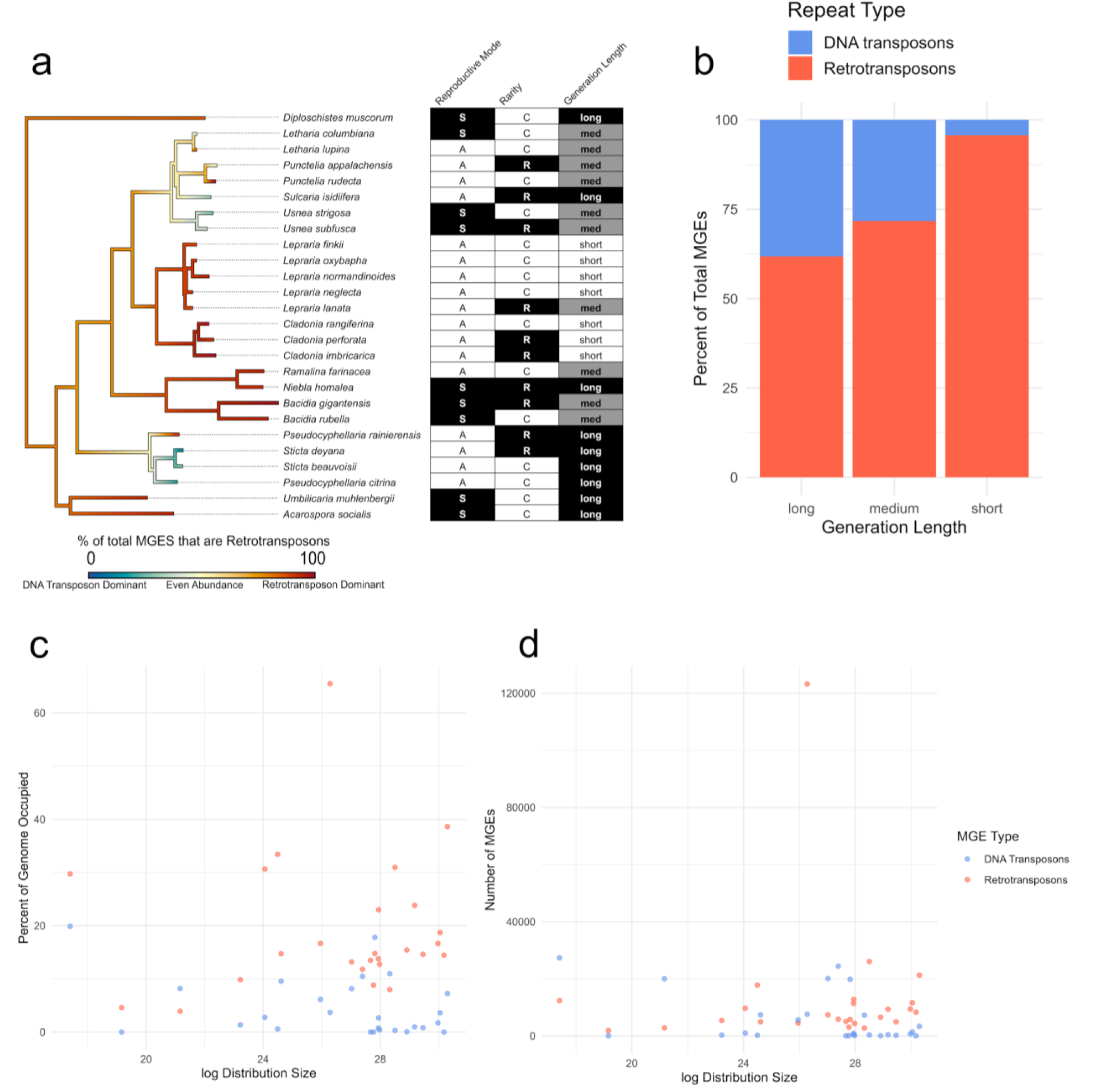
Relationship between MGE abundance and species traits. **a)** Species tree of all long-read genomes in the mobile genetic element (MGE) dataset, with RT abundance continuously mapped over the tree. Branches in red represent a genome with retrotransposon (RT) dominance, and branches in blue represent transposable element (TE) dominance. Branches in yellow have an even abundance of both RTs and TEs. The grid to the right of the phylogeny contains the data for reproductive mode (sexual (S) or asexual (A)), rarity (rare (R) or common (C)), and generation length for each species. **b)** Bar plot representing the mean proportion of all MGEs that were assigned as RTs (red) or TEs (blue) by generation length categories. **c)** Scatter plot of MGE abundance values for each species, with percent of the genome occupied on the y-axis and log of distribution size on the x-axis. Blue dots represent TE abundance, and red dots represent RT abundance. **d)** The same scatterplot, this time representing the number of MGEs in each genome on the y-axis.

Trait data were assembled from various sources for all species (Figure 2a; Table 2). Distribution sizes ranged from 3.62e7 meters squared to 1.44e13 meters squared, with rare species averaging a distribution of 1.71e11 (SE= 3.41e10) meters squared and common ones 5.25e12 (SE=1.24e12; Table 2). The dataset contained nine sexually reproducing species and 17 asexually reproducing species. There were 10 rare species and 16 common species. Nine of the species had a long generation length, ten had a medium generation length, and seven had a short generation length.

### Genomic MGE abundance is associated with traits of rarity in lichenized fungi

We recovered a significant relationship between the percent of the genome occupied by DNA transposons and log distribution size (PGLMM Binomial, p-value < 0.004; Figure 2d; Table 4), but did not find a significant relationship between RTs and log distribution size (Figure 2c) or total MGEs and log distribution size (Table 4). The percentage of the genome occupied by total MGEs was significantly higher in species with long generation lengths when compared to species with medium generation lengths (PGLMM Binomial, p-value < 0.004; Table 4). Further, species with long generation lengths had a significantly higher percentage of their genome occupied by DNA transposons than those with medium generation lengths (PGLMM Binomial, p-value < 0.004; Table 4). The number of total MGEs and DNA transposons in the genomes of species with long generation lengths was significantly higher than in those with medium or short generation lengths (PGLMM Poisson, p-value < 0.004; Table 5). For both rarity and reproductive mode, no significant relationship was recovered between either trait and the percent of the genome occupied by MGEs or the number of MGEs in the genome (Table 4-5).

**Table 4.**
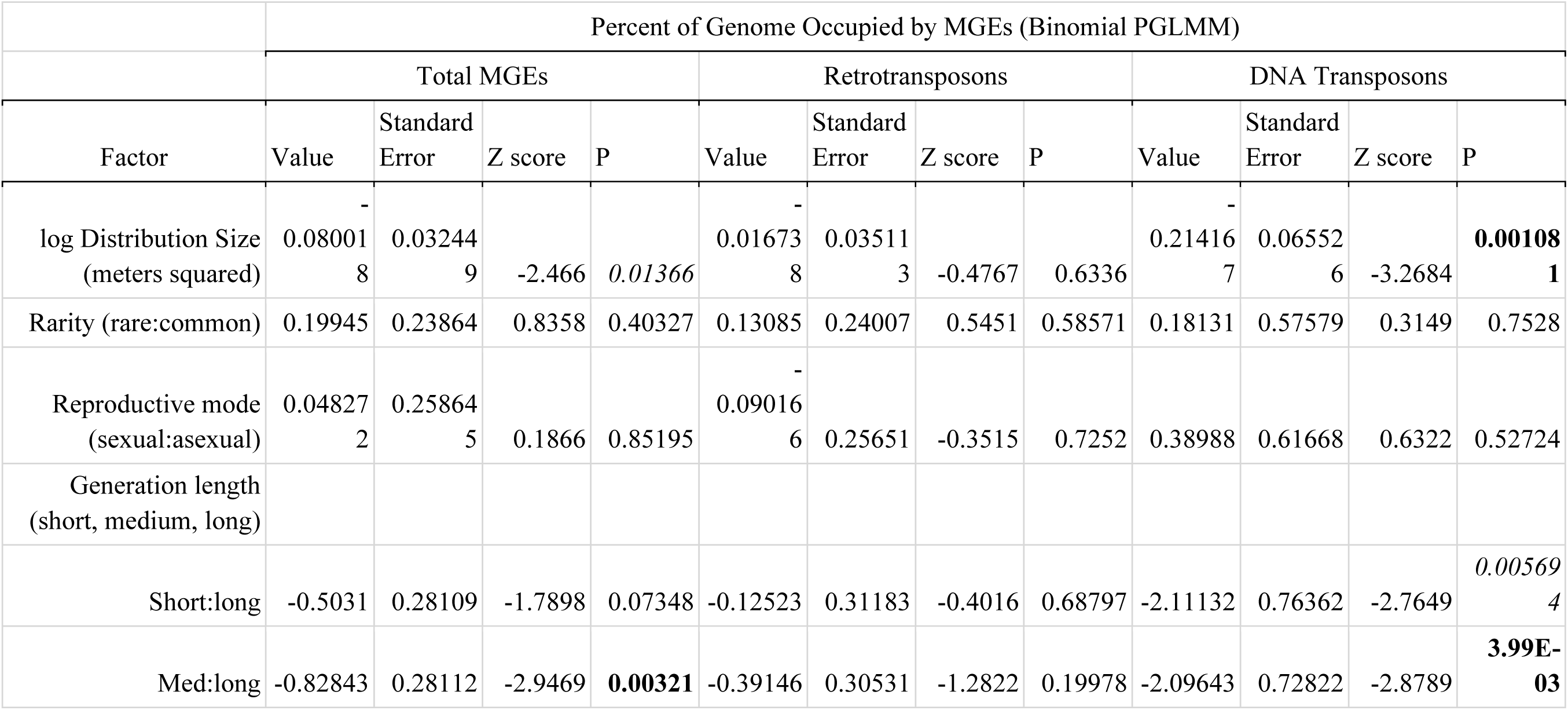
Binomial Phylogenetic Generalized Linear Mixed Model (PGLMM) Results and Test Values. Bolded values are significant with a p-value threshold corrected for multiple testing (Bonferroni, p-value < 0.004). Italicized p-values are less than 0.05.

**Table 5.**
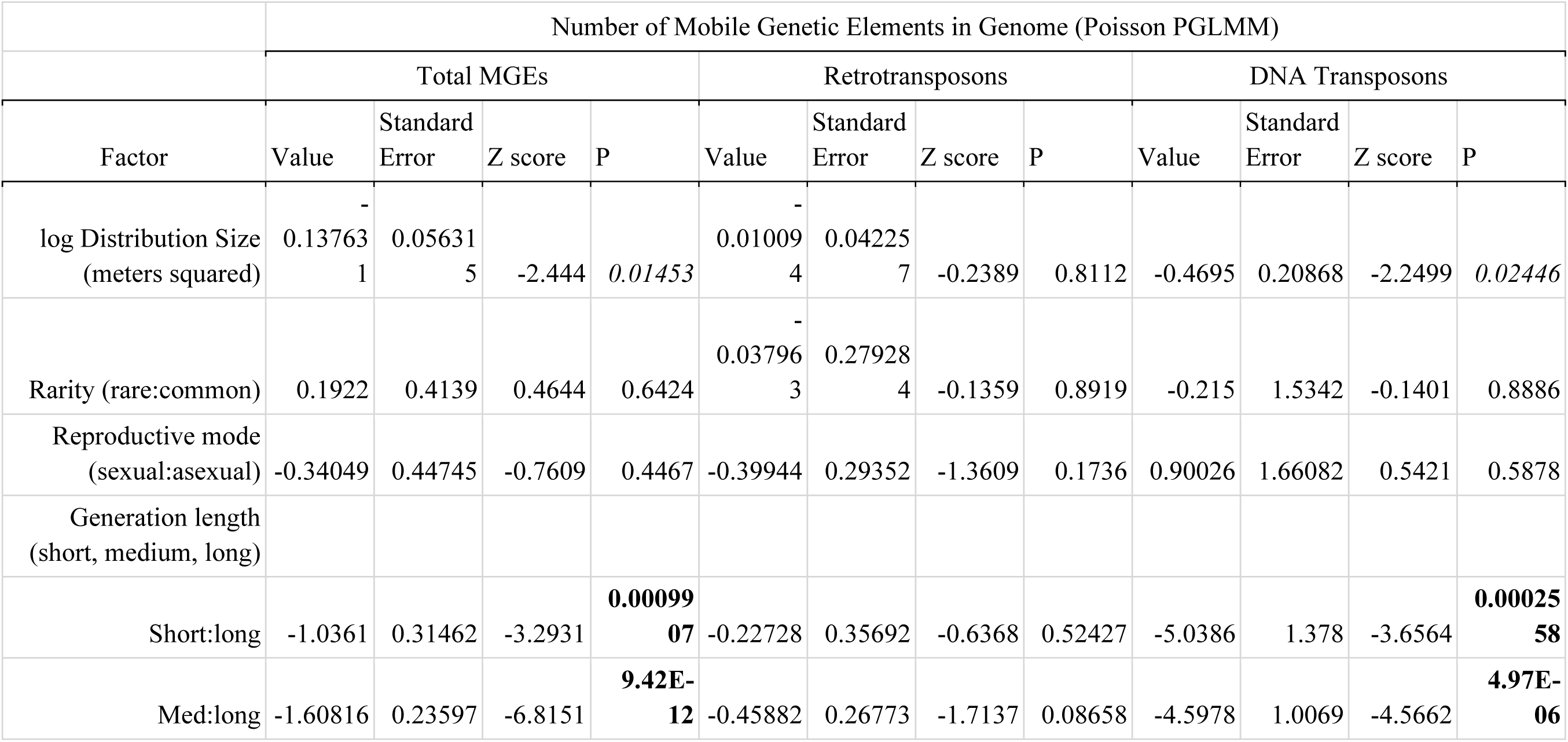
Poisson Phylogenetic Generalized Linear Mixed Model (PGLMM) Results and Test Values. Bolded values are significant with a p-value threshold corrected for multiple testing (Bonferroni, p-value < 0.004). Italicized p-values are less than 0.05.

|  | Number of Mobile Genetic Elements in Genome (Poisson PGLMM) |  |  |  |  |  |  |  |  |  |  |  |
| --- | --- | --- | --- | --- | --- | --- | --- | --- | --- | --- | --- | --- |
|  | Total MGEs |  |  |  | Retrotransposons |  |  |  | DNA Transposons |  |  |  |
| Factor | Value | Standard Error | Z score | P | Value | Standard Error | Z score | P | Value | Standard Error | Z score | P |
| log Distribution Size (meters squared) | -0.137631 | 0.056315 | -2.444 | *0.01453* | -0.010094 | 0.042257 | -0.2389 | 0.8112 | -0.4695 | 0.20868 | -2.2499 | *0.02446* |
| Rarity (rare:common) | 0.1922 | 0.4139 | 0.4644 | 0.6424 | -0.037963 | 0.279284 | -0.1359 | 0.8919 | -0.215 | 1.5342 | -0.1401 | 0.8886 |
| Reproductive mode (sexual:asexual) | -0.34049 | 0.44745 | -0.7609 | 0.4467 | -0.39944 | 0.29352 | -1.3609 | 0.1736 | 0.90026 | 1.66082 | 0.5421 | 0.5878 |
| Generation length (short, medium, long) |  |  |  |  |  |  |  |  |  |  |  |  |
| Short:long | -1.0361 | 0.31462 | -3.2931 | **0.0009907** | -0.22728 | 0.35692 | -0.6368 | 0.52427 | -5.0386 | 1.378 | -3.6564 | **0.0002558** |
| Med:long | -1.60816 | 0.23597 | -6.8151 | **9.42E-12** | -0.45882 | 0.26773 | -1.7137 | 0.08658 | -4.5978 | 1.0069 | -4.5662 | **4.97E-06** |

We assessed if either RTs or TEs were enriched in rare species by calculating the proportion of RTs relative to all MGEs in each genome, wherein all MGEs that were not classified as RTs were classified as TEs. We found a significant positive association between log distribution size and percent of all MGEs identified as RTs, as distribution size increases, RTs increase and, conversely, DNA transposons decrease. Thus, a greater proportion of MGEs are TEs in species with smaller distributions. Additionally, we found that species with long generation lengths were significantly enriched with DNA transposons when compared to species with medium and short generation lengths, which had more RTs (PGLMM Binomial, p-value < 0.0125; Figure 2b; Table 6). Because the largest number of DNA transposons in most species were attributed to the category of Rolling Circles (i.e., Helitrons and Unknown RCs; Supplementary Tables 3-5), we also tested if the relative abundance of Rolling Circles versus all other TEs was driving these relationships, and found again that Helitrons were enriched in species with small distribution sizes and long generation lengths (Figure 2b).

**Table 6.**
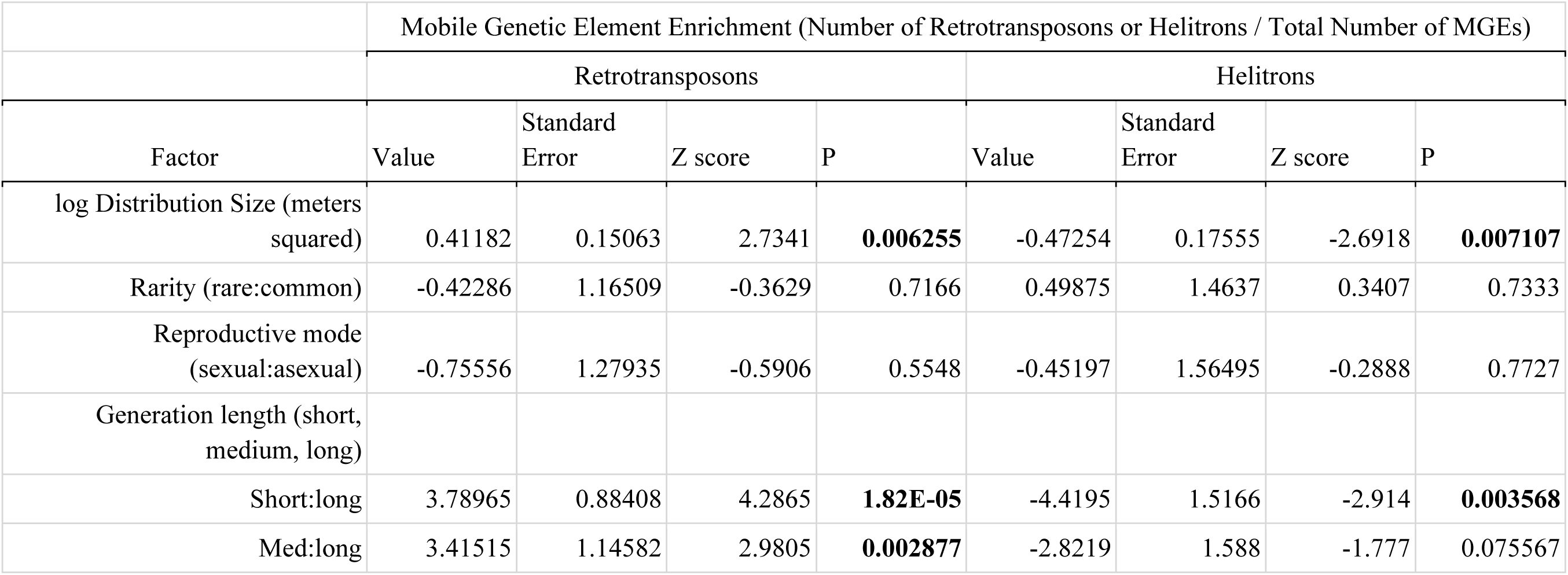
Mobile Genetic Element (MGE) Enrichment Binomial Phylogenetic Generalized Linear Mixed Model Results and Test Values. Bolded values are significant with a p-value threshold corrected for multiple testing (Bonferroni, p-value < 0.0125). Italicized p-values are less than 0.05.

|  | Mobile Genetic Element Enrichment (Number of Retrotransposons or Helitrons / Total Number of MGEs) |  |  |  |  |  |  |  |
| --- | --- | --- | --- | --- | --- | --- | --- | --- |
|  | Retrotransposons |  |  |  | Helitrons |  |  |  |
| Factor | Value | Standard Error | Z score | P | Value | Standard Error | Z score | P |
| log Distribution Size (meters squared) | 0.41182 | 0.15063 | 2.7341 | **0.006255** | -0.47254 | 0.17555 | -2.6918 | **0.007107** |
| Rarity (rare:common) | -0.42286 | 1.16509 | -0.3629 | 0.7166 | 0.49875 | 1.4637 | 0.3407 | 0.7333 |
| Reproductive mode (sexual:asexual) | -0.75556 | 1.27935 | -0.5906 | 0.5548 | -0.45197 | 1.56495 | -0.2888 | 0.7727 |
| Generation length (short, medium, long) |  |  |  |  |  |  |  |  |
| Short:long | 3.78965 | 0.88408 | 4.2865 | **1.82E-05** | -4.4195 | 1.5166 | -2.914 | **0.003568** |
| Med:long | 3.41515 | 1.14582 | 2.9805 | **0.002877** | -2.8219 | 1.588 | -1.777 | 0.075567 |

No statistically significant relationships were detected between MGE proximity to genes and trait data after applying a Bonferroni correction (adjusted p-value threshold = 0.00625; Gaussian PGLMM; Supplementary Table 6). However, possible trends were observed for longer generation length and asexual reproduction (Figure 1b and c, respectively), which were associated with decreased MGE distance to genes (p < 0.05), particularly for retrotransposons and DNA transposons in models without transcriptional directionality. The two *Pseudocyphellaria rainierensis* individuals collected from different populations high sequence similarity, including a similar number of MGEs (Figure 1b; Supplementary Table 3-4), and a pairwise alignment of the two genomes only revealed one small inversion on contig 4 (Figure 1d).

## DISCUSSION

### Genome metrics and architecture

We generated 18 new long-read genomes for Lecanoromycete lichenized fungi, contributing substantially to the availability of genomic resources for a group of organisms that is underrepresented in scientific research. Most genome’s metrics were in line with previously published long-read lichenized fungal genomes, with BUSCO scores greater than 90% and the size of most genomes ranging from 30 - 50 Megabases (Table 1). We noticed that *Pseudocyphellaria rainierensis* (131 Mb) and *Sulcaria isidiifera* (71 Mb) had much larger genomes than the rest. Upon annotating these genomes, it became clear that this size difference was due to high MGE content, as *P. rainierensis* and *S. isidiifera* had as much as 69% and 50% of their genomes occupied by MGEs, respectively, while most of the other genomes ranged from 10 to 30%. Sequencing a second *P. rainierensis* individual confirmed that the unusually high MGE, and especially RT content, was potentially a trait of the species genome and not specific to an individual population. *Pseudocyphellaria rainierensis* and *S. isidiifera* are both geographically restricted species, endemic to declining habitats, which led us to hypothesize that MGE abundance may be connected to rarity in lichenized fungi. Assembling the repetitive regions of short-read genomes is difficult without a long-read reference (Kellogg 2015). The completion of these new genomes from long read sequencing therefore provided us with a unique opportunity to investigate MGEs and genome architecture in lichenized fungi.

### DNA transposons, not retrotransposons, are associated with small distribution sizes

We hypothesized that RTs would have the strongest association with rarity because they have been linked to environmental stress and disturbances (Milyaeva et al. 2023; Miousse et al. 2015). Some RTs even contain promoter elements that allow them to transpose in response to specific stress stimuli. For example, Cavrak et al. (2014) found that *onsen* long terminal repeat (LTR) copia-like elements in *Arabidopsis thaliana* contain a promoter that activates their transposition as a response to plant heat shock transcription factors. Further, high RT content has been reported from some rare species. Linscott et al. (2022) analyzed the genome of a calcareous substrate endemic land snail, *Oreohelix idahoensis*, and found a high abundance of LTR RTs relative to other land snail genomes, with evidence suggesting recent expansions in LTR families. However, they did not identify any patterns related to DNA transposons. A study of the endangered fern *Vandenboschia speciosa* by Ruiz-Ruano et al. (2021) also recovered a large quantity of LTR RTs in the species’s genome, constituting 51% of the genome as opposed to the 5.6% of the genome occupied by TEs. Similarly, RTs were dominant in most of the genomes in our analysis when compared to TEs (Figure 3a, b), a phenomenon consistent with data from other fungal genomes (Castanera et al. 2017). We also found a significant negative association between MGEs and log distribution size, such that species with smaller distribution sizes had a higher percentage of their genome occupied by MGEs (Table 4). However, our statistical results showed that RTs were not the primary driving force of the patterns observed, and instead the relationship between MGEs and distribution size was primarily driven by TE abundance.

**Fig 3.**
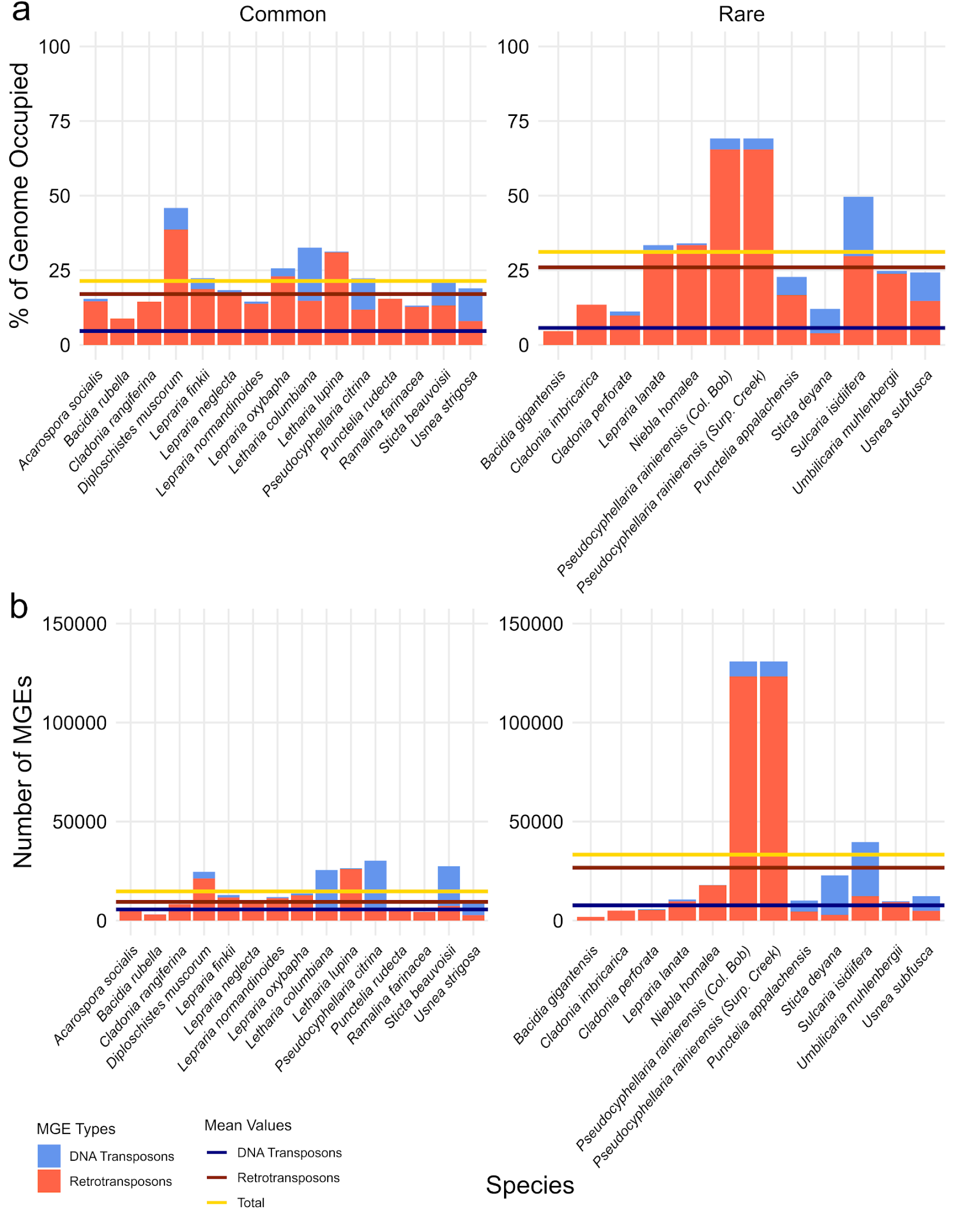
Overall Genomic MGE Results. **a)** Percent of the whole genome occupied by mobile genetic elements (MGEs) per species, with retrotransposons represented in red (bottom bar) and DNA transposons represented in blue (top bar). The left grid represents all species assigned as common in the dataset, with the right grid containing the rare species. Horizontal lines indicate the mean value for rare or common species for each MGE type: DNA transposons (dark blue), retrotransposons (dark red), and total MGE content (yellow). **b)** Number of MGEs in each species genome.

### Helitrons are significantly enriched in lichenized fungi with small distribution sizes

We found that the genomes of species with smaller distribution sizes and those with longer generation lengths were significantly enriched in DNA transposons—specifically, Helitrons and unknown Rolling Circles (Figure 1b, Table 6). Rolling Circles are one of the few TEs whose mechanism of movement enables them to self-replicate (Barro-Trastoy and Köhler 2024). Thus, while our hypothesis that RTs would be significantly enriched in rare species was not supported, our findings may still be consistent with our predicted mechanism of unchecked self-replication of MGEs in stressful environmental conditions. Helitrons have been previously identified in the genome of *Umbilicaria pustulata* where they comprise the majority of the Class II elements identified (Dal Grande et al. 2021). Notably, approximately 10% of the Helitrons found by Dal Grande et al. (2021) were highly differentiated insertions between populations from different ecotypes, potentially associating them with environmental stress. We expected to recover intraspecific variation in the high MGE content of *P. rainierensis*, but did not observe a difference in the abundance of MGEs of any type in the genomes of two individuals sequenced from different populations (Supplementary Tables 3-5), nor did we observe any impacts on the structure of the genome which could be caused by active MGEs (Figure 1d). Given that *P. rainierensis* grows in a narrow range of ecological conditions and in old growth habitat that is presumably very stable, it may not be a good model for understanding intraspecific MGE variation triggered by environmental influence, like what was observed in Dal Grande et al. (2021).

Helitrons are unique among TEs in that they have the ability to ‘capture’ and express host genes and regulatory elements (Barbaglia et al. 2011). This interaction poses a conflict for host MGE defenses, as silencing of Helitrons results in the silencing of captured host genes. A study by Muyle et al. (2020) analyzed the genome of Maize to identify Helitrons with captured host genes and their regulatory markings. They found that, on average, Helitrons with captured host genes were less impacted by host silencing when compared to other Helitrons. Additionally, Helitron captured host genes were more impacted by host silencing when compared to other host genes not captured by a Helitron (Muyle et al. 2020). Helitrons have variable transcriptional activity among species of Fungi, and data from a study by Castanera et al. (2014) suggested that they may be less transcriptionally active in Ascomycota than Basidiomycota. All taxa investigated here are Ascomycota suggesting that general patterns in Helitron activity may vary within phyla, but transcriptional data are required to test this hypothesis. The interactions among modification of gene expression through the activity of Helitron transposition and silencing, host-gene cargo, and rarity remains to be investigated.

### Generation length impacts MGE abundance

Generation length is a complex trait related to both life-span and reproduction. In lichens, generation length is primarily driven by lifespan with a minor influence of reproductive mode (Yahr et al. 2024); both sexually and asexually reproducing species may exhibit shorter or longer generation lengths, and, indeed, reproductive mode is not correlated with generation length in our dataset (Supplementary Table 2). Sexual reproduction may be predicted to either facilitate the spread of MGEs to new populations, or reduce MGE abundance by increasing purifying selection (Arkhipova and Meselson 2004; Bestor 1999). Testing these competing hypotheses, Nowell et al. (2021) found that in Bdelloid rotifers, which reproduce exclusively asexually, TEs evolved in a primarily random fashion, with no apparent relationships between reproductive mode and MGE content being recovered. We similarly did not find any significant relationships between MGE abundance and reproductive mode, but a marginal trend (p < 0.05) was uncovered for MGE spatial distribution, indicating that asexually reproducing species may experience higher instances of MGE and gene colocalization (Supplementary Table 6). In termites, MGE transcription profiles are found to cluster based on both age group and reproductive caste (Post et al. 2022). Previous studies have found surprisingly low rates of somatic mutation and high genome integrity in plants and fungi with long life spans (Hofmeister et al. 2020; Anderson et al. 2018; Schmid-Siegert et al. 2017). In our study, species with long generation lengths had a higher number of MGEs, specifically TEs, in their genomes than those with short or medium generation lengths, contradicting these previous findings. The potential interactions among life-span, reproductive mode, generation length, and MGE dynamics in the context of complex lichen life cycles warrants further investigation.

### Ploidy, life cycles, and MGEs

Lichenized fungi are commonly assumed to consist of haploid mycelia, which only enters a brief dikaryotic diploid phase upon the formation of ascomata during sexual reproduction (Tripp and Lendemer 2018). Thus, species with long generation lengths, or those who form only haploid asexual spores and propagules, spend the vast majority of their time in a haploid state.

Hiltunen et al. (2022) suggest that higher MGE activity is associated with haploidy. They compared the expression level of MGEs in different phases of the life cycle of the fairy-ring mushroom, *Marasmius oreades*, and found that TE mutagenesis was most active during the haploid phase of the life cycle, and the formation of dikaryotic hyphae after sex and fusion resulted in a highly stable genome with little TE activity. If MGE activity is associated with a haploid genome state (Hiltunen et al. 2022), haploid species may be the most at risk for MGE accumulation and resultant genome mutations. As the available genomic resources for lichenized fungi increases, it has been revealed that the assumption that lichen’s vegetative hyphae are always formed by haploid mycelia is not true, which may be an additional factor contributing to MGE variation (Tripp et al. 2017; McKenzie et al. 2020). In our study, all of the genomes originated from haploid thalli except that of *Letharia lupina*, which is triploid, and *Letharia columbiana*, which is diploid (McKenzie et al. 2020), thus this hypothesis remains to be tested.

Bryophytes are another group of organisms with a haploid dominant life-cycle. They also share a poikilohydric physiology with lichens. The MGE landscape in bryophyte genomes is not well characterized, but it is known that variations in genome size among species are not fully explained by whole genome duplication events (Szövényi et al. 2021). A study by Patel et al. (2025) found that hornworts had the smallest genomes with the least size variation when compared to liverworts and mosses. Variability of MGE abundance has been observed in several groups of non-vascular land plants, and MGE activity could be one possible explanation for size variation among this group (Kirbis et al. 2025; Linde et al. 2023; Lang et al. 2018). Due to the alternation of sexual and asexual reproductive modes in the bryophyte life cycle, increased focus on generating genomic resources for bryophyte MGEs may provide further insights into the intersecting relationship between generation length, reproductive mode, and MGE activity in haploid dominant organisms using an evolutionarily independent comparison point.

## CONCLUSION

Previous studies of single rare species have recovered large abundances of RTs occupying their genomes resulting in a focus on RTs and their association with rarity and genomic stress in conservation research (Ruiz-Ruano et al. 2021; Linscott et al. 2020; Abascal et al. 2016). We similarly found two rare species with extraordinarily high RT content in their genomes. However, by investigating MGE abundance in a multi-species comparative phylogenetic framework, we uncovered unexpected associations between distribution size, generation length, and TE abundance in lichenized fungi. Specifically, TEs, and especially self-replicating Helitrons, were significantly enriched in species with small distribution sizes and long generation lengths while RTs were not. Further research is needed to determine if these patterns are present in other groups of organisms, which will require continued focus on the development of high-quality, long-read reference genomes for more species. Assessing rarity for effective species conservation requires a multifaceted approach. MGEs, particularly TEs, may serve as a genomic signature of rarity, providing additional context for conservation management and strengthening our understanding of the processes that increase extinction risk.

## ACKNOWLEDGEMENTS

Funding for this project was provided by NSF DEB #2115191, Start-up funds from Eastern Washington University, and support from the ORG.one program to JLA, NSF DEB #2436848 to JCL, and the Stuntz Mycology Fund, Puget Sound Mycological Society Ben Woo Research Grant, Washington Native Plant Society Research and Plant Inventory Grant (#23-EN-RPI-02), American Bryological and Lichenological Society Culberson & Hale Grant for Field Research in Lichenology, and The Evergreen State College Summer Undergraduate Research Fellowship to STS and LC. Field and lab work by Eli Balderas, as well as field work conducted by Jason Hollinger and Bert Harris were essential for completing this project.

## DATA AVAILABILITY

Sequence data are deposited at the National Center for Biotechnology Information (BioProject PRJNA1270085). Code is available at https://github.com/jallen73/Lichen-mobile-elements.

